# *De novo* mutations in domestic cat are consistent with an effect of reproductive longevity on both the rate and spectrum of mutations

**DOI:** 10.1101/2021.04.06.438608

**Authors:** Richard J. Wang, Muthuswamy Raveendran, R. Alan Harris, William J. Murphy, Leslie A. Lyons, Jeffrey Rogers, Matthew W. Hahn

**Affiliations:** Department of Biology, Indiana University, Bloomington, IN; Human Genome Sequencing Center, Baylor College of Medicine, Houston, TX; Department of Molecular and Human Genetics, Baylor College of Medicine, Houston, TX; Veterinary Integrative Biosciences, Texas A&M University, College Station, TX; Department of Veterinary Medicine and Surgery, College of Veterinary Medicine, University of Missouri, Columbia, MO; Department of Computer Science, Indiana University, Bloomington, IN

**Author notes:** Corresponding author: Richard J. Wang.

## Abstract

The mutation rate is a fundamental evolutionary parameter with direct and appreciable effects on the health and function of individuals. Here, we examine this important parameter in the domestic cat, a beloved companion animal as well as a valuable biomedical model. We estimate a mutation rate of 0.86 × 10^-8^ per bp per generation for the domestic cat (at an average parental age of 3.8 years). We find evidence for a significant paternal age effect, with more mutations transmitted by older sires. Our analyses suggest that the cat and the human have accrued similar numbers of mutations in the germline before reaching sexual maturity. The per-generation mutation rate in the cat is 28% lower than what has been observed in humans, but is consistent with the shorter generation time in the cat. Using a model of reproductive longevity, which takes into account differences in the reproductive age and time to sexual maturity, we are able to explain much of the difference in per-generation rates between species. We further apply our reproductive longevity model in a novel analysis of mutation spectra and find that the spectrum for the cat resembles the human mutation spectrum at a younger age of reproduction. Together, these results implicate changes in life-history as a driver of mutation rate evolution between species. As the first direct observation of the paternal age effect outside of rodents and primates, our results also suggest a phenomenon that may be universal among mammals.

## Introduction

Mutation is a fundamental force in evolution, the primary source of genetic novelty, and often a fitness burden to its bearers in the form of inherited disease. Germline mutations, and the processes that govern the occurrence and rate of these heritable genetic changes, have long been of interest to evolutionary biologists (e.g. Muller 1928; Crow 2000). Approaches for studying the mutation rate have changed drastically with changes in sequencing technologies. The oldest estimates of mutation rates predate the availability of molecular data and were based on the incidence of rare dominant disease (Haldane 1935; reviewed in Nachman 2004). With low-cost whole-genome sequencing, direct estimates of the mutation rate can now be made by comparison of parents and offspring. These estimates of the per-generation mutation rate in a growing number of species have enabled an increased understanding of how this fundamental parameter evolves.

A generation is the natural unit for expressing a mutation rate when we measure the number of mutations an individual inherits from its parents. However, since generation times vary among species, per-generation estimates of the mutation rate across species are difficult to compare. Nevertheless, current data suggest at least a twofold range of mutation rates among primates (Chintalapati and Moorjani 2020). In almost all primate species studied to date, paternal age has an effect on the number of inherited mutations (Kong et al. 2012; Venn et al. 2014; Goldmann et al. 2016; Thomas et al. 2018; Besenbacher et al. 2019; Wang et al. 2020; Bergeron et al. 2021). Because males transmit so many more mutations to their offspring than females, the paternal age effect dominates in comparisons of the total number of inherited mutations.

Together, the paternal age effect and male mutation bias obscure the mechanisms driving mutation rate evolution: have per-generation mutation rates changed due to differences in the quality of DNA repair or are they due to changes in life-history? Variation in the age of reproduction across species, coupled with parental age effects (Rahbari et al. 2016; Jónsson et al. 2017), would suggest that per-generation rates are expected to vary across species simply because of differences in life-history.

One approach that attempts to appropriately compare mutation rates between species is to take the calendar age of the parents at conception into account (e.g. Besenbacher et al. 2019).

Given a known relationship between parental age and the number of mutations inherited by offspring, this approach can predict differences in the mutation rate between species solely due to differences in the age at reproduction for parents from each species. Deviations from this prediction imply changes apart from life-history, be they cellular, molecular, or environmental. This approach assumes that mutations accumulate at a constant rate across the lifespan of an individual (see also Gao et al. 2019), and is therefore equivalent to comparing per-year estimates of the mutation rate (Besenbacher et al. 2019; Chintalapati and Moorjani 2020). However, germline mutations appear to accumulate at different rates across different life stages (Rahbari et al. 2016; Scally 2016; Harland et al. 2017; Jónsson et al. 2018; Sasani et al. 2019; Jonsson et al. 2021). Furthermore, the estimated relationship between parental age and the number of inherited mutations inferred from trio studies can only be predictive to the age of puberty: the number of germline mutations cannot be directly observed via transmission before parents reach reproductive age. Therefore, a model of “total longevity” (i.e., calendar age) in comparisons between species may not fully account for differences in mutation accumulation across different life stages. This is especially true when the age at puberty differs between species.

Differences in the rate of mutation accumulation across life stages have motivated an alternative model that splits the process into two regimes: before and after the onset of sexual maturity (e.g. Thomas and Hahn 2014; Amster and Sella 2016; Gao et al. 2016). The disproportionate contribution of mutations from fathers—and the biology of spermatogenesis, which requires continuous division of the male germline after puberty (Crow 2000)—strongly suggest that reproductive age marks an important change in the rate of mutation accumulation. A starting point for comparisons between species is therefore to take the reproductive age of parents into account. Like the total longevity model described above, this “reproductive longevity” model (Thomas et al. 2018) assumes a common relationship among species between parental age and the number of mutations inherited by offspring. The key distinction for this model is that age-related mutation accumulation only begins with the onset of puberty; the number of mutations before sexual maturity is assumed to be the same between species, and independent of the age at which puberty is reached. In this model, differences in the per-generation mutation rate between species are best explained by reproductive longevity: that is, the length of time for mutation accumulation after puberty. Even when the per-generation mutation rate has evolved between species, this model can help to delineate the stage(s) at which changes have occurred (Wang et al. 2020).

To date, most pedigree studies of mutation have been focused on humans and other primates (e.g., Kong et al. 2012; Venn et al. 2014; Besenbacher et al. 2019, but see Lindsay et al. 2019 for an estimate in mouse). To better understand how mutation rates evolve, a broader collection of species will need to be considered. Distinguishing between models for the evolution of mutation rates will also require species with a larger span of generation times and greater differences in the age of puberty. In this study, we present a direct estimate of the mutation rate from the domestic cat (*Felis catus*). Cats are an important companion animal for many households and a common target for interventions by veterinary medicine. Recent studies of genetic variation in the cat and the development of feline genomic tools (e.g., Genova et al. 2018; Buckley et al. 2020; Bredemeyer et al. 2021) have advanced the domestic cat as a biomedical model. Reliable mutation rates would also help to accurately estimate the evolutionary history of domesticated and wild cat species (Figueiró et al. 2017; Li et al. 2019).

In this study, we consider models for the evolution of mutation rates in the domestic cat. The difference in lifespan between cats and longer-lived primates, like humans, provide a strong contrast for testing models of mutation accumulation. In addition to considering a model of mutation rates across species, we perform a novel analysis contrasting the mutation spectrum across species. As with the total number of inherited mutations, the mutation spectrum also changes with the age of parents (Jónsson et al. 2017). Therefore, differences in mutation spectrum between species with very different ages at reproduction may also be explained by changes in life-histories. Our results suggest that a model of reproductive longevity explains most of the difference in both the rate and spectrum of mutations between the cat and the human.

## Results

### Mutation rate in the domestic cat

We sequenced individuals from 11 domestic cat trios to a median of 41× coverage using Illumina short-read sequencing. Of the 22 total individuals sequenced, 19 were part of a larger pedigree, while 3 were from a standalone trio (Fig. 1). DNA from this standalone trio was collected from cultured fibroblasts, while all others were collected from blood samples. After filtering sites based on mapping quality and coverage, we retained an average 1.8 Gb out of the 2.5 Gb reference genome, from which we identified candidate *de novo* mutations in each trio. Briefly, we considered sites on autosomes where both parents were homozygous for the reference allele and the offspring was heterozygous for an alternate allele not observed elsewhere in the dataset (see Methods). We applied a set of stringent filters to the initial list of candidates and found 233 single nucleotide *de novo* mutations across the 11 trios (Supplemental Data 1). The structure of the pedigree allowed us to trace mutations to a third generation in four grandoffspring, and we found transmission in 40 out of the 78 potential instances, close to the expected 50% transmission rate. We combined haplotype-sharing information for mutations from these families with read-pair phasing at all mutations to determine the parent of origin for 124 mutations. We found 93 mutations were of paternal origin while 31 were of maternal origin, consistent with a male mutation bias (binomial *p* = 2.2 × 10^-8^; Wilson Sayres and Makova 2011; Ségurel et al. 2014). The proportion of male-biased cat mutations (75%) is consistent with the proportion found in humans (80.4%; Jónsson et al. 2017) and other mammals (de Manuel et al. 2022). These phased mutations also show a parental age effect: more mutations were transmitted from older parents, significantly so for older fathers (Poisson regression of read-phased mutations, paternal *p* = 0.032; Fig. S1).

**Figure 1.**
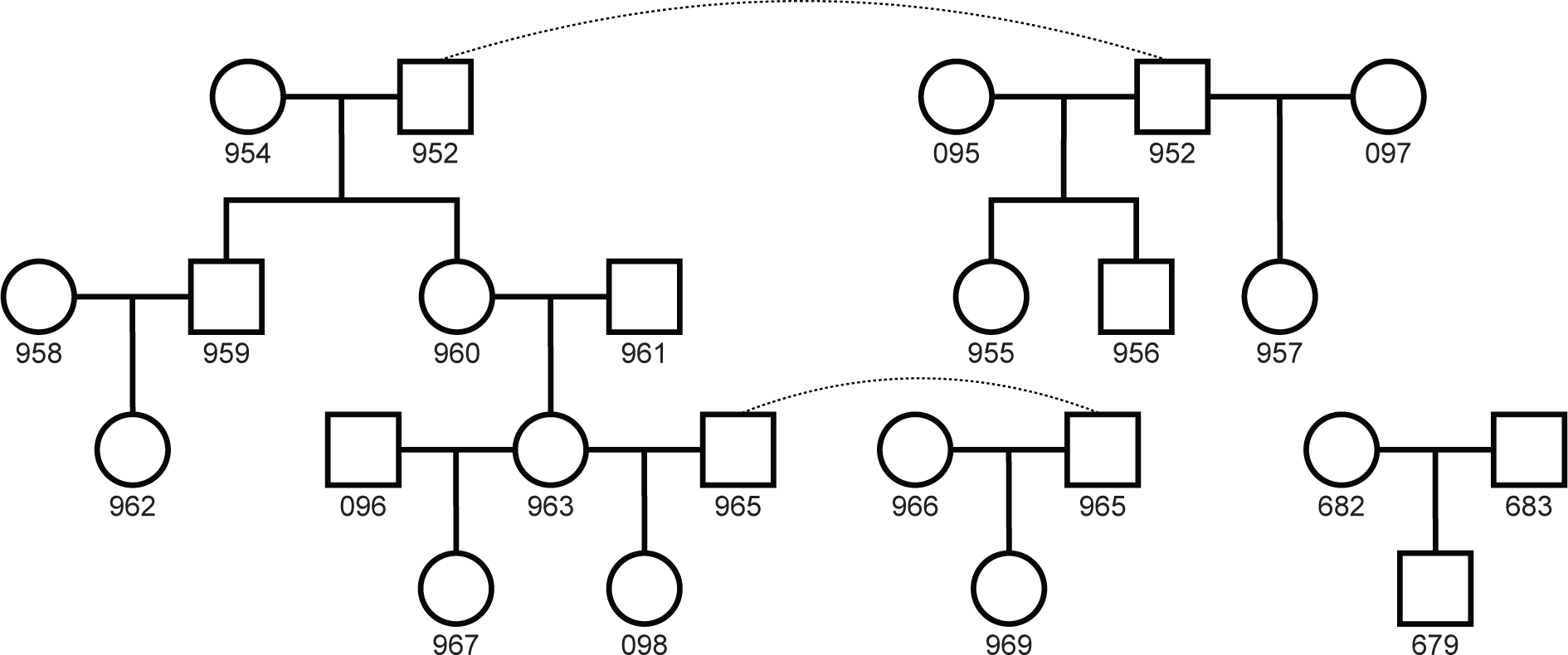
Pedigree of sequenced individuals The 22 total individuals sequenced form 11 separate trios for mutation analysis. Whole blood was sampled from the 19 individuals in the larger pedigree, while cultured fibroblasts were sampled from the 3 individuals in the standalone trio. Dashed lines connect identical individuals in the pedigree.

To estimate the per-generation mutation rate, we divided the number of mutations identified in each trio by the observed genome size and corrected for the false negative rate from our stringent set of filters. We estimated this latter value as the “site callability,” by calculating the fraction of sites that pass our filters from a sample of sites across the genome in each trio (see Methods; Besenbacher et al. 2019; Wang et al. 2020). This approach produced stable estimates of the mutation rate as the filter stringency was increased (Fig. S2), providing confidence in our estimate. We found the mean per-generation mutation rate in cat to be 0.86 × 10^-8^ [95% CI: 0.75, 0.97] per bp for parents at an average age of 3.8 years across sexes (♂: 4.7 y, ♀: 2.9 y), compared to 1.29 × 10^-8^ per bp in humans with an average age of 30.1 years (Jónsson et al. 2017); Table 1 shows the rate estimated for each cat trio separately. Notably, the mutation rate estimated for the trio using DNA isolated from cultured fibroblasts was not notably different from those using DNA from blood samples.

**Table 1.** Mutation counts and the per generation mutation rate

| Proband ID<sup>1</sup> | Age at conception (y) | | Mutations | Mean depth<sup>2</sup> | Callability | Haploid Size (Mb) | Rate ($\times 10^{-8}$) |
| --- | --- | --- | --- | --- | --- | --- | --- |
|  | Paternal | Maternal |  |  |  |  |  |
| 679<sup>3</sup> | - | - | 22 | 47.8 | 0.644 | 1493 | 1.14 |
| 955 | 3.9 | 4.4 | 21 | 38.8 | 0.714 | 1655 | 0.89 |
| 956 | 3.9 | 4.4 | 7 | 37.0 | 0.620 | 1611 | 0.35 |
| 957 | 12.0 | 1.4 | 29 | 42.7 | 0.751 | 1674 | 1.15 |
| 959 | 4.8 | 3.3 | 20 | 39.5 | 0.671 | 1609 | 0.93 |
| 960 | 4.8 | 3.3 | 24 | 38.2 | 0.695 | 1643 | 1.05 |
| 962 | 3.4 | 2.1 | 20 | 42.0 | 0.723 | 1880 | 0.74 |
| 963 | 2.3 | 2.8 | 17 | 40.6 | 0.711 | 1923 | 0.62 |
| 967 | 1.7 | 2.7 | 17 | 39.4 | 0.719 | 2070 | 0.57 |
| 969 | 3.9 | 1.6 | 27 | 39.7 | 0.676 | 1952 | 1.02 |
| 098 | 6.1 | 3.1 | 29 | 42.5 | 0.729 | 2018 | 0.99 |

<sup>1</sup>ID of offspring for each independent trio in Figure 1
<sup>2</sup>Mean read depth across individuals of the trio with offspring as proband
<sup>3</sup>Sequences from this trio were derived from cell culture.

Assuming the average parental age in our sample (3.8 years) is representative of the average age of reproduction in the cat, we estimate a per-year mutation rate of 2.3 × 10^-9^ [2.0, 2.5] per bp. This is much higher than the human rate of 0.43 × 10^-9^ per bp per year (Jónsson et al. 2017), or the reported per-year rate from any primate (Besenbacher et al. 2019; Wang et al. 2020; Wu et al. 2020; Campbell et al. 2021). The higher rate in cats is driven by the similar number of mutations at puberty, but a much shorter generation time. We also calculated the per-year substitution rate from sequence divergence in unconstrained regions between *F. catus* and *Panthera tigris* (Zoonomia Consortium 2020), which diverged approximately 12 million years ago (Figueiró et al. 2017). This comparison resulted in an estimate of 1.54 × 10^-9^ [1.2, 2.1] per year per bp, where the lower and upper bounds represent substitution rates calculated using the 95% highest posterior density interval for divergence times between these two species (Figueiró et al. 2017). The overlap between pedigree-based and phylogeny-based mutation estimates (assuming substitution rates reflect mutation rates), implies that the mutation process has been relatively unchanged over the last 12 million years of feline evolution.

The mutation spectrum in the cat largely resembles the spectrum found in the human and most other primates (Fig. S3). The largest difference that is significant was a lower percentage of A>G transitions in the cat, 19%, compared to the 27% found in humans (χ^2^ test, *p* < 0.005; human data from Jónsson et al. 2017). This was offset by a slightly higher percentage of C>T transitions, 48% in cat vs. 42% in human, leading to an overall transition / transversion ratio of 1.99 in cat. As in humans, C>T transitions at CpG sites accounted for a substantial fraction of all observed mutations (21%). We estimate the mutation rate at CpG sites in the cat to be 1.76 × 10^-7^ [1.26, 2.26] per bp per generation. Note that the median GC content in the reference genomes for the cat and the human are not substantially different (41.9% cat, 40.4% human; NCBI). We compared the mutation spectrum in the cat to the spectrum calculated from low-frequency variants segregating in a published cat dataset (Buckley et al. 2020). We found significant differences in the proportion of A>G and A>T mutations between the two (Fig. S3). Interestingly, a similar comparison of the human mutation and rare variant spectrum revealed significant differences as well (see Supplemental Results).

### Testing the reproductive longevity model of the mutation rate

Direct estimates of the mutation rate have consistently demonstrated a strong parental age effect in primates, with more mutations inherited by the offspring of older parents. To consider whether such an effect exists in the domestic cat, we performed a Poisson regression on the total number of mutations (adjusted by observable genome size) with parental age, and found a significant effect of paternal age on the mutation rate (*p* = 0.008). Each additional year of paternal age at conception leads to an estimated 3.1 (95% CI: [0.8, 5.4]) total additional mutations in the offspring. This estimate for the effect size of paternal age on the total number of mutations is not significantly different from those observed in humans (unequal variances *t*-test on data from Jónsson et al. 2017, *p* = 0.22). Using only the phased mutations, we find a significant paternal age effect that translates to an estimated 1.7 [0.2, 3.3] additional paternal mutations per year (*cf*. 1.51 per year of additional paternal age in humans; Jónsson et al. 2017). While we found no significant effect of maternal age on the mutation rate (Fig. S1), the effect we estimated was in the positive direction; much larger sample sizes were also used to detect a significant maternal age effect in humans (Goldmann et al. 2016; Jónsson et al. 2017).

The accumulation of mutations with age post-puberty among species is modeled in both the total longevity and reproductive longevity models (Fig. 2a). The key difference between the two models is in the number of mutations at sexual maturity. In the total longevity model, the cat mutation rate at sexual maturity is assumed to be an extension of the regression fit, extending the parental age effect found in humans to the age at sexual maturity in the cat. In contrast, the reproductive longevity model includes a parameter for the number of mutations that occur before puberty in each species, with the parental age effect only acting post-puberty (Fig. 2a). We compared the predictions made by these two models to the data from our cat trios: Figure 2b shows the increase in per-generation mutation rates with years of paternal age after puberty (assuming a puberty age of 0.5 y in cat, 13 y in human; Tsutsui et al. 2004; Parent et al. 2003).

**Figure 2.**
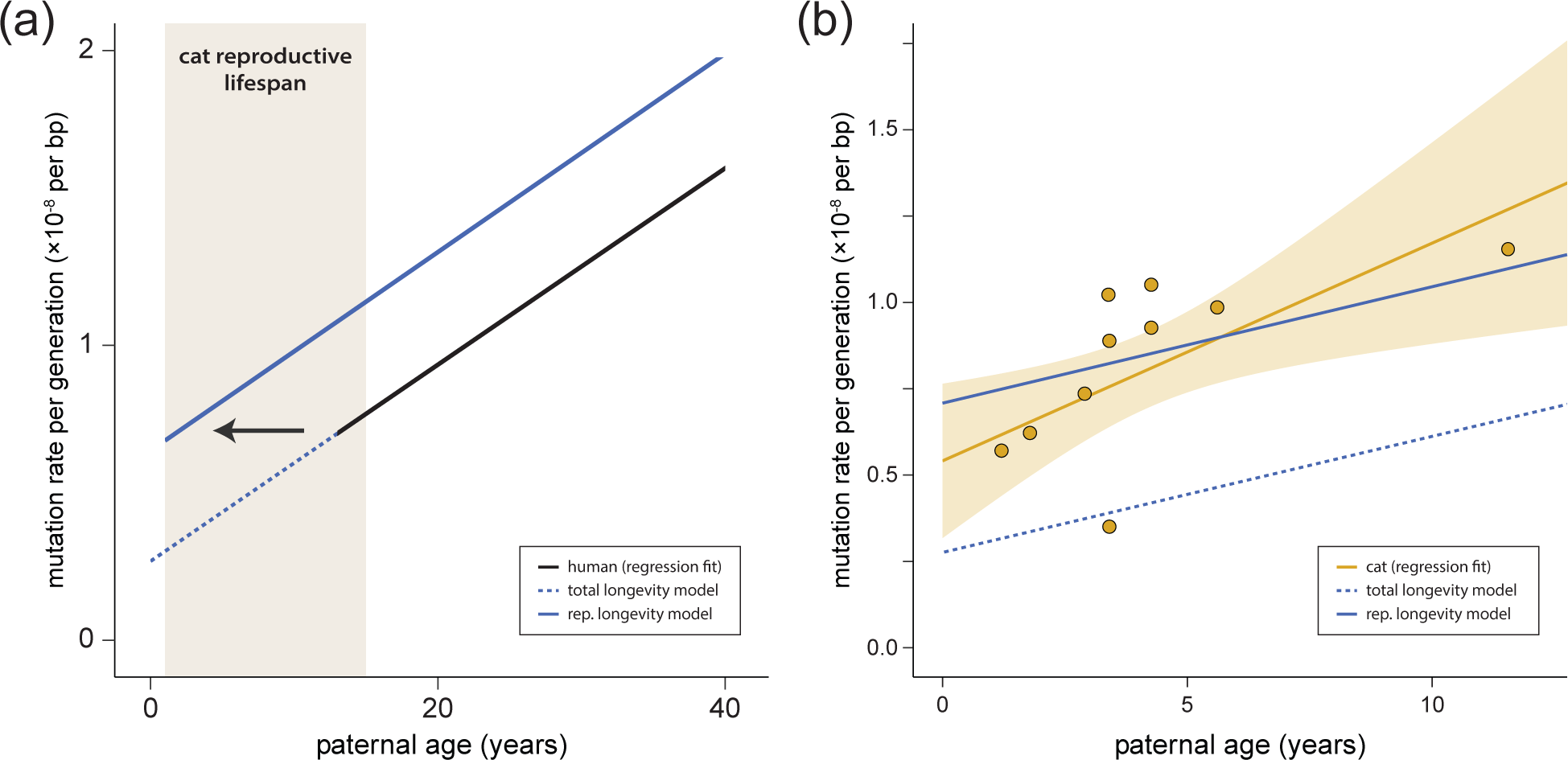
Mutation rates increase with paternal age (a) The predicted per-generation mutation rate in the domestic cat based on human data (regression fit) using the reproductive longevity model and the total longevity model. (b) The per-generation mutation rate increases with paternal age in the domestic cat; each point represents mutation data from one trio. Fit line shows regression under a Poisson model with the 95% CI shaded. The reproductive longevity model provides a much better fit to the observed pattern with age (*t*-test of residuals; *p* = 0.0002).

The reproductive longevity model provides a significantly better fit to the mutation rates observed in the cat (*t*-test of residuals, *p* = 0.0002).

Using the regression fit presented in Figure 2b, we found no significant difference between cat and human in per-generation mutation rates estimated at the age of puberty (*t*-test, *p* = 0.41): we estimate 0.59 × 10^-8^ per bp (95% CI: [0.39, 0.78]) in cat vs. 0.67 × 10^-8^ per bp ([0.65, 0.69]) in human (Jónsson et al. 2017). While this simple regression fit may miss biological phenomena in the unobserved period before sexual reproduction, the consistency of the estimated rate between species who shared a common ancestor approximately 85 million years ago (Tarver et al. 2016) suggests to us a conserved pre-puberty developmental program among mammals, as has previously been noted in owl monkeys (Thomas et al. 2018). Note that the number of pre-puberty mutations was slightly lower in rhesus macaques (Wang et al. 2020).

These results do not explain how estimates of male-mutation bias appear to be relatively constant post-puberty (see Gao et al. 2019), suggesting some additional process that generates this bias before puberty.

### A reproductive longevity model of the mutation spectrum

Parental age affects not only the number of *de novo* mutations inherited by offspring, but also their composition (Carlson et al. 2020). In humans, the proportion of C>T transitions decreases with paternal age while the proportion of C>G transversions increases with maternal age (Goldmann et al. 2016; Jónsson et al. 2017). The dependence of the mutation spectrum on age suggests multiple underlying mutation processes whose relationships differ with age (Goldmann et al. 2019). Since differences in life history appear to explain much of the variation in per-generation mutation rates between cat and human, we were interested in knowing whether a model of reproductive longevity could be used to explain differences in the mutation spectrum. That is, does the mutation spectrum in the cat resemble the mutation spectrum of a human that has had the same amount of post-pubertal time for mutation accumulation? Or, alternatively, are there different mutational processes in cats that explain the difference in mutation spectrum?

To build a reproductive longevity model of the mutation spectrum, we used mutation data from a large human dataset (Jónsson et al. 2017). We modeled the accumulation of mutations for each mutation class as independent and fit the number of mutations in each class with a Poisson regression. From this regression, we predicted the spectrum for a given age by dividing the predicted number of mutations in each class by the total number of mutations predicted across classes. Figure 3a shows how the predicted mutation spectrum changes with parental age relative to the average spectrum observed in the human dataset (with an average age of 30.1; Jónsson et al. 2017). Under this model, the mutation spectrum for species with different parental ages at conception can vary dramatically from the one observed in humans without any change to the mutational process between species.

**Figure 3.**
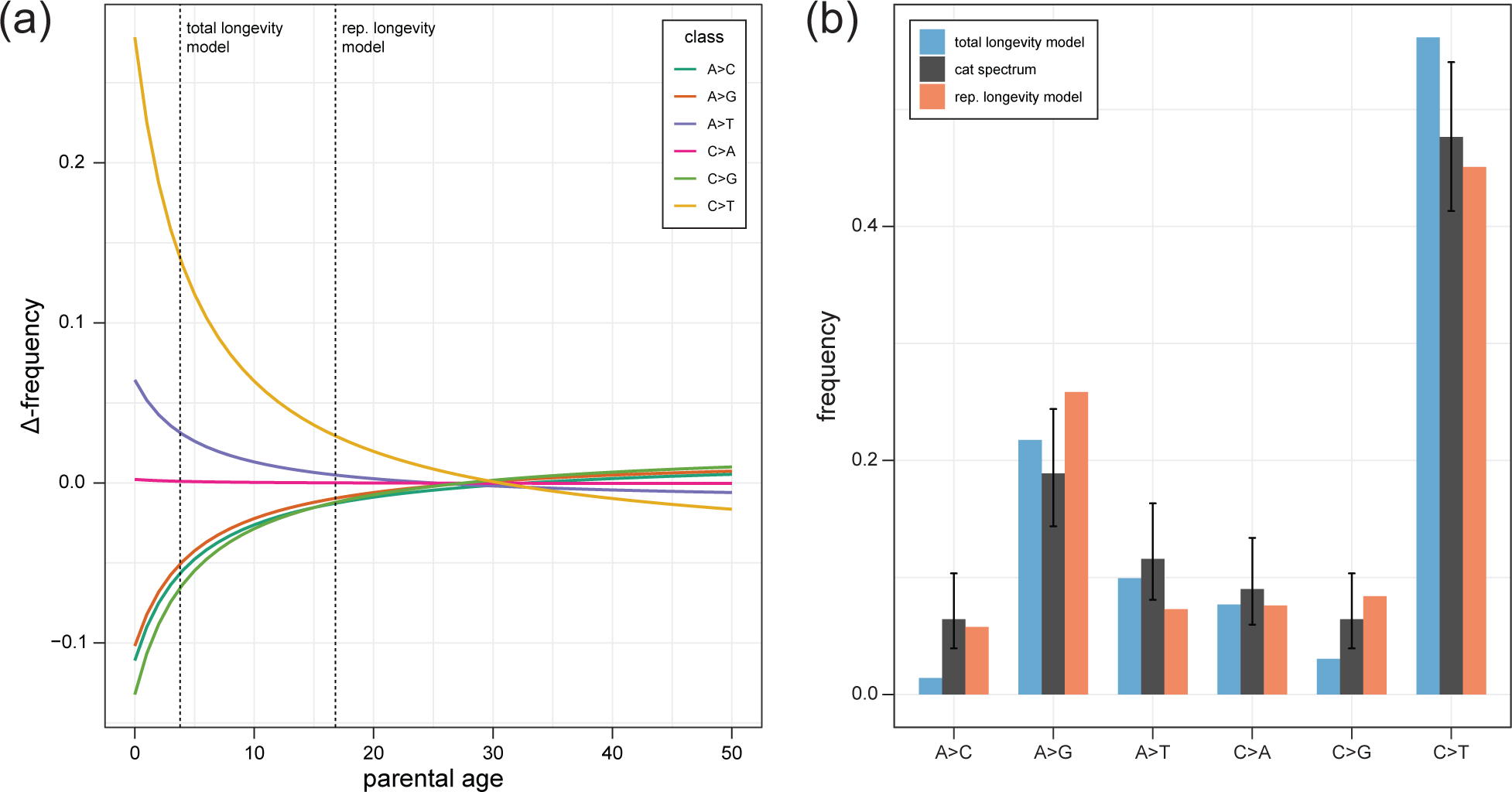
Longevity models of the mutation spectrum (a) The predicted change in each mutation class as a function of average parental age (colored lines). The predictions come from fitting observed human data using a Poisson regression (see main text). The vertical lines indicate the spectrum that would be predicted under a total longevity model and a reproductive longevity model. (b) The observed cat mutation spectrum (black bars) compared to the predicted frequency of each mutation class under a total longevity model and a reproductive longevity model. Error bars on the cat spectrum show a binomial 95% CI (Wilson score interval) for their respective classes.

We compared the fit of the observed cat mutation spectrum (at an average parental age of 3.8 y) to the predicted spectrum under both the total longevity and reproductive longevity models. The predicted mutation spectrum for the cat is expected to depart more from the average human spectrum under the total longevity model than under the reproductive longevity model because there is a larger difference in the ages considered (calendar age vs. reproductive age; Fig. 3a). We found that the predicted spectrum under the reproductive longevity model provided a better fit to the observed cat mutation spectrum (Fig. 3b; root mean square error 3.6% [95% CI: 3.2, 4.3] vs. 4.5% [1.7, 7.4]). Under the better-fitting reproductive longevity model, the predicted proportion of C>T transitions, 45.1%, closely matched the one found in the cat spectrum, 47.6%.

The most notable difference between this prediction and the observed spectrum was the proportion of A>G transitions, which accounted for 25.9% of mutations in the prediction, but only 18.8% of observed mutations in the cat. While the overall fit of the spectrum predicted by the total longevity model was worse, its prediction for the proportion of A>G transitions and A>T transversions was much closer to the observed spectrum in the cat (predicted vs. observed for A>G and A>T respectively: 21.7% vs. 18.9% and 9.9% vs. 11.6%). Together, these results suggest that, while the reproductive longevity model captures much of the mutation spectrum’s dependence on age, multiple mutation processes likely underlie differences in mutation spectra between species.

## Discussion

We find a mutation rate and spectrum for the domestic cat that is variable throughout its lifespan and consistent with a model of reproductive longevity. The estimated per-generation rate, at 0.86 × 10^-8^ per bp, is lower than that found in the human but, because of the shorter lifespan of the cat and similar number of mutations pre-puberty between species, corresponds to a higher per-year rate. Our results are among the first to demonstrate the existence of a paternal age effect from direct estimates outside of primates (Figure 2b). In conjunction with direct estimates in mice (Lindsay et al. 2019), our results suggest that the paternal age effect on mutation rate may be universal to all mammals. Although power to detect a paternal age effect may differ among studies, an effect of similar magnitude has been found in almost every mammalian species examined (Kong et al. 2012; Venn et al. 2014; Goldmann et al. 2016; Thomas et al. 2018; Besenbacher et al. 2019; Lindsay et al. 2019; Wang et al. 2020; Bergeron et al. 2021; see Wu et al. 2020 for an exception in the baboon). Differences in the magnitude of the parental age effect are not the only way in which the mutation rate could evolve. Changes to the number of mutations before reaching puberty, as suggested in macaques (Wang et al. 2020), will affect the per-generation rate without any change in the parental age effect. However, we found no significant difference in the number of mutations before puberty between cats and humans (Figure 2b).

Pedigree studies in primates have underscored the difference between mutation rates measured within species and substitution rates measured between species (Chintalapati and Moorjani 2020). In humans, the difference between these two rates has called into question assumptions about how mutation rates translate into substitution rates (Scally and Durbin 2012; Ségurel et al. 2014; Thomas and Hahn 2014; Amster and Sella 2016; Gao et al. 2016).

Connecting the two will require a better understanding of how mutation rates vary across a lifespan, especially pre- and post-puberty. This dichotomy is the major difference between the total longevity and reproductive longevity models (Figure 2a). Determining whether rates of mutation accumulation or the number of mutations at puberty differ in mammals is difficult, as most data have come from closely related primates (e.g. Besenbacher et al. 2019) that have highly similar ages at puberty and lifespans. Estimates of the mutation rate in mice (Lindsay et al. 2019), which have an even shorter generation time than cats, show a similar contrast to the one we have presented between cats and humans, and are consistent with the reproductive longevity hypothesis. The per-generation estimate in mice is lower than in cats (0.39 × 10^-8^ vs. 0.86 × 10^-8^ per bp) while the per-year estimate is higher (5.3 × 10^-9^ vs. 2.3 × 10^-9^ per bp).

Interestingly, our estimate of the per-year substitution rate between cats (1.5 × 10^-9^ per bp) has confidence intervals that overlap with our pedigree-based estimate of the mutation rate, in contrast to the discrepancies observed in primates. However, our estimate of the per-generation mutation rate in the cat is nearly double the estimate in the wolf (*Canis lupus*), another member of the order Carnivora, at 0.45 × 10^-8^ per bp (Koch et al. 2019). Given the relatively similar age at reproduction for the wolves used in that study (mean: 3.1 y), and the consistency of our estimates with a reproductive longevity model and with phylogenetic estimates of the per-year rate, we cannot say where the discrepancy in these estimates lies. Nevertheless, distinguishing between modes of evolution in the mutation rate will require data from more species with varied life histories. Figure S4 shows a comparison of the per-generation mutation rate estimated in the cat relative to other mammals.

The spectrum of mutations in the cat was remarkably similar to those found in most primates, with a preponderance of C>T and A>G transitions (Kong et al. 2012; Venn et al. 2014; Thomas et al. 2018; Besenbacher et al. 2019; Wang et al. 2020; Wu et al. 2020; Bergeron et al. 2021; see Campbell et al. (2021) for an exception in the mouse lemur). We showed that differences in the mutation spectrum between the cat and the human can in large part be explained by differences in age according to a model of reproductive longevity. Our results indicate that, like mutation rate itself, the mutation spectrum should be treated as a function-valued trait that changes with parental age. Mutation spectra have typically been reported and compared with little regard for the reproductive age of individuals in the sample. But just as in the case for mutation rates, meaningful comparisons must take reproductive age and longevity into account. This applies not only to comparisons of spectra between species and populations (Harris and Pritchard 2017), but also to inferences about spectra from different stages of the germline (Rahbari et al. 2016; Bae et al. 2018). Distinct spectra from different developmental stages have been used to imply that a unique set of mutational processes underlie each stage (Goldmann et al. 2019). However, a single set of processes pervading all stages is equally capable of producing significant differences in spectra. Distinguishing between these competing hypotheses requires the comparison of models for mutation spectra that consider reproductive age.

In addition to sequencing blood from individuals in the larger pedigrees, we sequenced DNA taken from cell lines developed from a single trio. Researchers routinely sequence blood samples for studies of *de novo* mutations to avoid sampling a small number of cells from a single clonal lineage that may contain a somatic mutation at high frequency. Furthermore, cell lines that have been passaged for a long period can accumulate mutations at a very high rate: in the 1000 human genomes project (Altshuler et al. 2010), 90-95% of mutations initially detected were cell line-specific. Notably, the estimate here from a single trio of cultured fibroblasts was not distinguishable from the blood-derived estimates. The strategy for identifying and filtering mutations did not need to be different for the cell-line samples in our analysis, but the collected fibroblasts proceeded through only two or three passages before DNA was collected, which may have mitigated any error-prone behavior.

Finally, our results have important implications for multiple aspects of feline evolution and veterinary care. Accurate estimates of mutation rates are important for reconstructing the evolutionary history of cats. By combining genomic data from domesticated cats with data from the wildcat (*Felis silvestris lybica*), our mutation rate estimates will allow for more refined estimates of the timing of cat domestication (Driscoll et al. 2007; Ottoni et al. 2017) and felid evolution in general (Figueiró et al. 2017; Li et al. 2019). Cat breeding, and the management of captive felids, has altered population dynamics over the past few thousand years. In nature, the normal life expectancy for wild cat species has been reported to be between 2 and 5 years of age (Figueiró et al. 2017; Li et al. 2019). Both the breeding of domestic cat breeds and species survival plans of wild felids allow breeding in older animals, advancing the reproductive age for these populations. The results presented here suggest that advanced parental age may lead to the production of offspring with parental-age related maladies and diseases, as in humans (Hassold and Hunt 2001; Malaspina et al. 2002; Kong et al. 2012). Therefore, population managers may need to become more vigilant to the health problems in offspring associated with advanced paternal age.

## Methods

### Samples and sequencing

Archived whole blood samples were provided from 19 domestic cats in an extended pedigree taken from a cross-bred colony of cat breeds maintained for biomedical disease model characterization. The disease models did not compromise growth or reproductive maturity of the cats (Lyons et al. 2004; Lyons et al. 2016; Cogné et al. 2020). All animal care and use was conducted in accordance with policies and guidelines approved by the IACUC of the University of Missouri, protocol 8313. In addition, cultured fibroblasts from a trio of three domestic cats were isolated at the National Cancer Institute, Poolesville animal colony.

Genomic DNA was isolated from samples for whole genome sequencing. Standard PCR-free libraries were prepared using KAPA Hyper PCR-free library reagents (KAPA Biosystems). Total genomic DNA was sheared into fragments of approximately 200-600 bp and purified using AMPure XP beads. Sheared DNA molecules were subjected to double size selection with different ratios of AMPure XP beads. This was followed up with DNA end-repair and 3’- adenylation before the ligation of barcoded adapters. Library quality was evaluated by fragment analysis and qPCR assay. These WGS libraries were sequenced on an Illumina HiSeq X, producing 150 bp paired-end reads.

### Mapping and variant calling

Sequenced reads were aligned with BWA-MEM v. 0.7.12-r1039 (Li 2013) to the domestic cat reference genome Felis_catus_9.0 (Buckley et al. 2020) using the parameters “bwa mem -M -t 8” with other parameters being default. Picard MarkDuplicates v. 1.105 (Broad Institute 2019) was used to identify and mark duplicate reads from the BAM files. We used GATK version 4.1.2.0 (Van der Auwera et al. 2013) to call variants using best practices (https://gatk.broadinstitute.org/hc/en-us/articles/360035535932-Germline-short-variant-discovery-SNPs-Indels). HaplotypeCaller was used to generate gVCF files for each sample using the parameters “gatk HaplotypeCaller -R --emit-ref-confidence GVCF” with other parameters being default. GenomicsDBs were generated from the gVCFs using GenomicsDBImport with default parameters. Variant discovery was performed jointly across samples and then genotyped with GenotypeGVCFs using default parameters. GATK SelectVariants --select-type-to-include SNP was used to generate the SNP vcf. We applied GATK hard filters: (SNPs: “QD < 2.0 || FS > || MQ < 40.0 || MQRankSum < -12.5 || ReadPosRankSum < -8.0”) and removed calls that failed.

### Filters for candidate mutations

An initial set of candidate mutations was identified as Mendelian violations in each trio.

Specifically, we looked for violations where both parents were reference homozygous and the offspring was heterozygous for an alternate allele. As this is the most common type of genotyping error (Wang et al. 2021), we then applied the following stringent filters to this initial set of candidates to get a set of high-confidence candidates:

- Read depth at the candidate site must be between 20 and 60 for every individual in the trio. Sites with too few reads may be sampling errors while sites with too many reads may be problematic repetitive regions (Li 2014).
- High genotype quality (GATK) in all individuals, (GQ > 70).
- Candidate mutations must be present on both the forward and reverse strand in the offspring (ADF, ADR > 0).
- Candidate mutation must not be present in any reads from either parent (AD = 0).
- Candidate mutation must not be present in samples that are not descended from the parents of each trio.
- Candidate mutation must not have low allelic depth in the offspring (Allelic Balance > 0.35).

We assessed the sensitivity of our mutation rate estimates across a range of stringency criteria and found them to be in good agreement across reasonable filter limits (see Fig. S2). We applied stringent filters, as listed above, with the intent of limiting false discoveries.

### Estimating the per-generation mutation rate

To convert the raw number of candidate mutations into an estimate of the mutation rate, we applied existing strategies that considered differences in coverage and filtering (see (Besenbacher et al. 2019; Wang et al. 2020). In brief, the number of identified mutations was divided by the total callable sites: a product of the number of sites covered by the appropriate sequencing depth and the estimated probability that a site would be called correctly given that it was a true de novo mutation. The mutation rate is then calculated as:

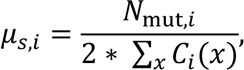

where μ_*s,i*_ is the per-base mutation rate for trio *i*, and *N*_mut,*i*_ is the number of *de novo* mutations in trio *i*, and *C*_*x*_ is the callability of site *x* in that trio. This strategy assumes the ability to call each individual in the trio correctly is independent, allowing us to estimate *C*_*x*_(*x*) as:

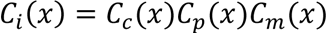

where *Cc*, *Cp*, and *Cm* are the probability of calling the child, father, and mother correctly for trio *i*. These values are estimated by applying the same set of stringent filters to high-confidence calls from each trio. For heterozygous variants in the child,

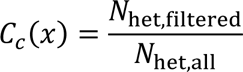

where *N*_het,all_ is the number of variants where one parent is homozygous reference and the other parent is homozygous alternate, leading to high confidence in the child heterozygote call, and *N*_het,filtered_ is the set of such calls that pass our child-specific candidate mutation filters. The parental callability, *C*_*p*_(*x*) and *C*_*m*_(*x*), were estimated in a similar manner, by calculating the proportion of remaining sites in each after the application of the stringent mutation filters. Finally, to calculate confidence intervals for reported mutation rates, we assumed a Poisson variance in the number of observed mutations.

*Phasing* de novo *mutations*

We determined the parent of origin for 45 mutations in probands 959, 962, and 963 by tracking transmission to a third generation. We accomplished this by comparing haplotype blocks in 50 kb regions surrounding each *de novo* mutation across individuals in the three-generation pedigrees. Mutations transmitted to a third generation were assigned to the parent matching the transmitted block while mutations that were not transmitted were inferred to be from the other parent. Haplotype blocks were established using genotypes at biallelic sites and the parent of origin was left undetermined if blocks were distinguished by fewer than two informative sites or appeared to have been subject to recombination (resulting in multiple genotype mismatches between blocks).

We used read-pair tracing to determine the parent of origin for 108 mutations across all of our trios. We did this by applying WhatsHap 1.0 (Patterson et al. 2015) in read-based phasing mode for each individual separately, and then matched informative blocks bearing the mutation to their parent of origin according to the rules of Mendelian inheritance. Ambiguous blocks, including any that showed genotype inconsistencies between parent and offspring, were left unphased. We combined the parent of origin inferred from both approaches, omitting one mutation that was inconsistent between the two methods, to arrive at the combined set of 124 phased mutations.

### Poisson regression

To estimate the effects of parental age on mutation rate, we modeled the number of mutations in a trio, *i*, with a Poisson regression as:

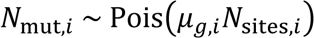

where *μ*g,*i* is the per-generation mutation rate for trio *i* and *N*_sites,*i*_ is the diploid callable genome size for trio *i*. We begin with an identity link Poisson regression:

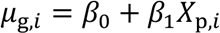

where *X*p,*i* is the paternal age for trio *i*, with *β0* and *β1* regression coefficients. To adjust for differences in the observable genome size, we generated a new variable:

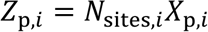

and fit the Poisson regression on *N*sites,*i* and *Z*p,*i* with no intercept. That is:

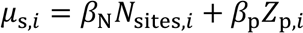

where *μ*s,*i* is the per-site per-generation mutation rate in trio *i*.

In this regression model, the product *β*_N_*N*_sites,*i*_ represents a per-site version of the original intercept, and the coefficient *β*p is the per-site scaled effect of paternal age on the per-generation mutation rate. We also performed this regression with an additional maternal age term, that is, *β*_m_*Z*_m,*i*_ where *Z*_m,*i*_ = *N*_sites,*i*_ *X*_m,*i*_ and *X*m,*i* is the maternal age for trio *i*, but removed it from the final analysis because the maternal coefficient was not significant in the regression.

Comparisons to the paternal age effect in humans applied the above regression framework to the full set of mutations identified in Jónsson et al. (2017).

For the read-phased *de novo* mutations featured in Figure S1, we performed regressions on the number of male and female mutations separately with a Poisson model on respective parental age using an identity link function. To estimate the paternal age effect from these phased mutations, we scaled the paternal regression coefficient by the fraction of mutations that were successfully read-phased.

### Reproductive longevity model of the mutation spectrum

We assume the relationship between parental age and the number of mutations in the offspring is independent for each parent (i.e. no interaction between maternal and paternal age).

Let *c* be the class for a given mutation, where *c* ∈ { A>C, A>G, A>T, C>A, C>G, C>T }. Let *Nc*,*i* be the number of mutations of class *c* in individual *i* from a given parent. We model

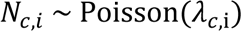

where λ*c*,*i* is the per-generation rate parameter for class *c* mutations in individual *i*, and is fit to a Poisson regression using an identity link with

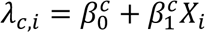

where *Xi* is the age at conception of the given parent for individual *i*. Let *N*^′^_*c*_ (*X*^′^) be the predicted number of mutations of class *c* at a given parental age *X*^′^. The predicted set of mutations from a parent with age *X*^′^ is then

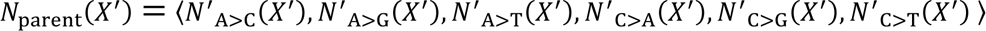

The spectrum for an individual with parents having ages *X’*mother and *X’*father can then be calculated as

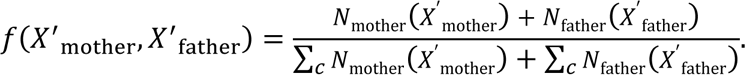

This model is initially fit with the mutations and parental ages from a large human dataset (Jónsson et al. 2017). From this fit, the cat mutation spectrum under the total longevity model was then predicted at the average parental age of individuals in our sample; i.e., as *f*(*X*^′^_mother_ = 2.92 years, *X*^′^_father_ = 4.68 years). Under the reproductive longevity model, we shifted the average parental age in our sample by the time to puberty in humans (13 years), predicting a cat mutation spectrum for our sample that resembles a human spectrum at an average age of 16.8 years (*X*^′^_mother_ = 15.92, *X*^′^_father_ = 17.68 years). To compare the two predictions, we calculated the root mean square error to the observed cat spectrum across all mutation classes. We calculated confidence intervals on the root mean square error by bootstrap resampling individuals from the human dataset. Coefficients in the regression for each mutation type were recalculated and used to predict spectra in 1,000 resampling replicates.

## Acknowledgements

This work was funded by the Precision Health Initiative of Indiana University with support for LAL from the University of Missouri, Gilbreath McLorn Endowment. We wish to thank Harsha Doddapaneni, Donna Muzny, and the sequence production team at the Human Genome Sequencing Center for support in data production and Nicole Foley at Texas A&M for sharing data. We also thank the associate editor and two reviewers for constructive comments.

## Data Availability

All sequencing data underlying this study has been submitted to the NCBI Sequence Read Archive (SRA) at https://www.ncbi.nlm.nih.gov/sra, and can be accessed under the BioProject with accession number: PRJNA740309. The software code for reproducing the figures from our supplementary data are freely available at: https://github.com/Wang-RJ/catMutations.

## Supplemental Material

### Supplemental Results

#### Comparisons of the mutation spectrum to the rare variant spectrum

We compared the cat mutation spectrum to the spectrum of rare polymorphisms from a published study of cat variation. To calculate the rare variant spectrum, we considered variants from the 99 Lives dataset (Buckley et al. 2020) at frequencies less than 5% in the folded frequency spectrum. We excluded singletons to reduce potential sequencing errors. Table S1 shows a comparison of this calculated spectrum to the mutation spectrum. A>G transitions are significantly overrepresented while A>T transversions are significantly underrepresented (both *p* < 0.05) in the rare variant spectrum relative to the mutation spectrum.

We performed the same comparison in humans, using extremely rare variants from Carlson et al. (2018) to calculate the polymorphism spectrum and mutations from Jónsson et al. (2017) to calculate the mutation spectrum. There is more power to detect differences with the larger number of mutations in the human dataset. We find that all but the proportion of A>T transversions are significantly different at the *p* < 0.05 level between the rare variant spectrum and the mutation spectrum for humans (Table S1).

**Table S1.** Comparison of mutation and variant spectra in cat and human

| cat spectrum | mutation class (percent) |  |  |  |  |  |
| --- | --- | --- | --- | --- | --- | --- |
|  | A>C | A>G | A>T | C>A | C>G | C>T |
| mutation | 6.44 | 18.88 | 11.59 | 9.01 | 6.44 | 47.64 |
| rare variant | 7.38 | 27.31 | 5.70 | 8.54 | 7.48 | 43.59 |
| human spectrum |  |  |  |  |  |  |
| mutation | 7.04 | 26.81 | 6.80 | 7.59 | 9.60 | 42.15 |
| rare variant | 7.30 | 27.23 | 6.98 | 10.19 | 8.81 | 39.49 |

## Supplemental Figures

**Figure S1.**
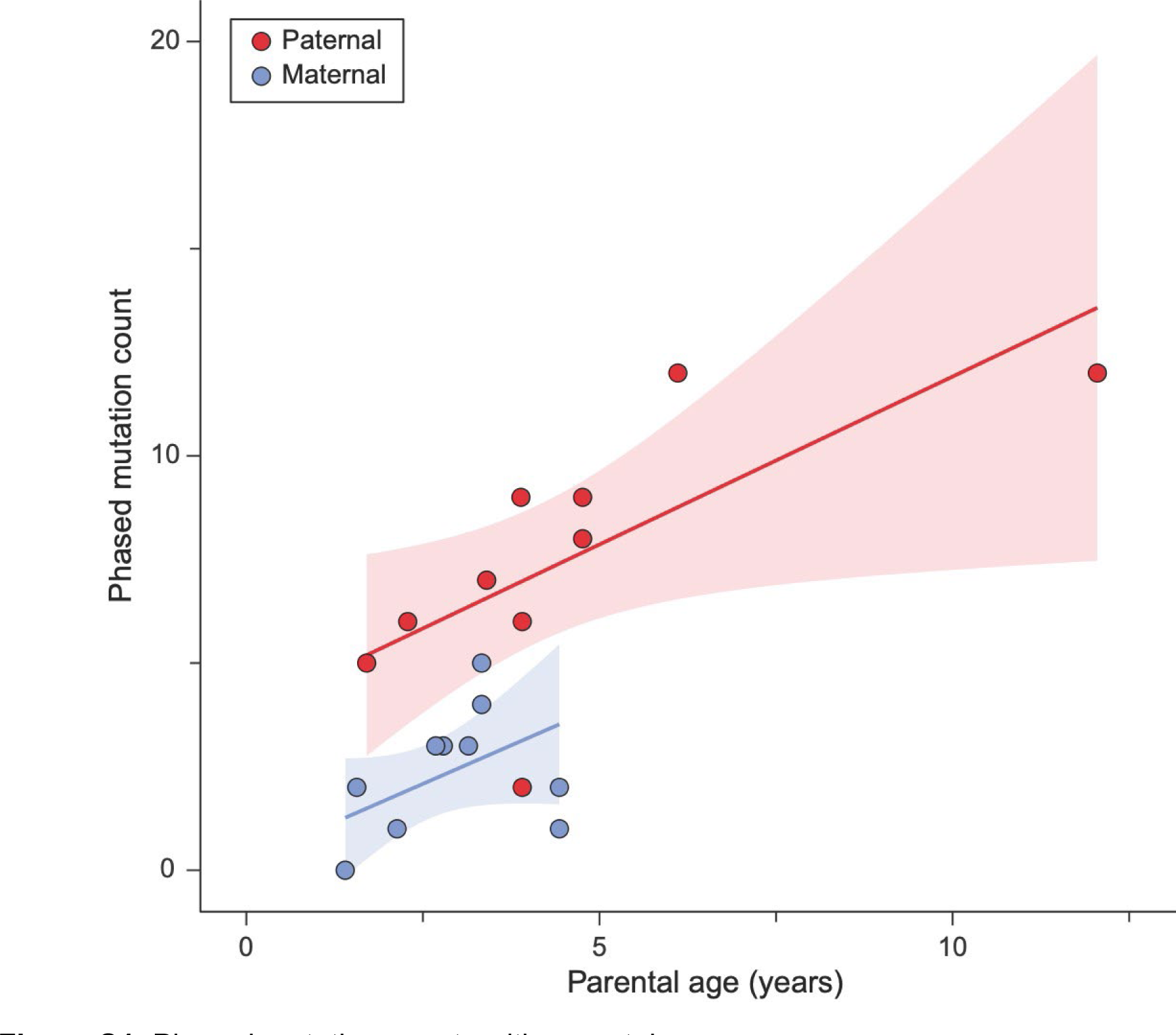
Phased mutation counts with parental age The number of phased mutations from ten trios of the domestic cat with a maternal (blue) and paternal origin (red), as identified by read-pair phasing. There is a positive relationship between the number of phased mutations and increasing parental age, though fewer maternal mutations limit the significance of its p-value (Poisson regression, paternal *p* = 0.032, maternal *p* = 0.116). Shaded areas show regression 95% CI.

**Figure S2.**
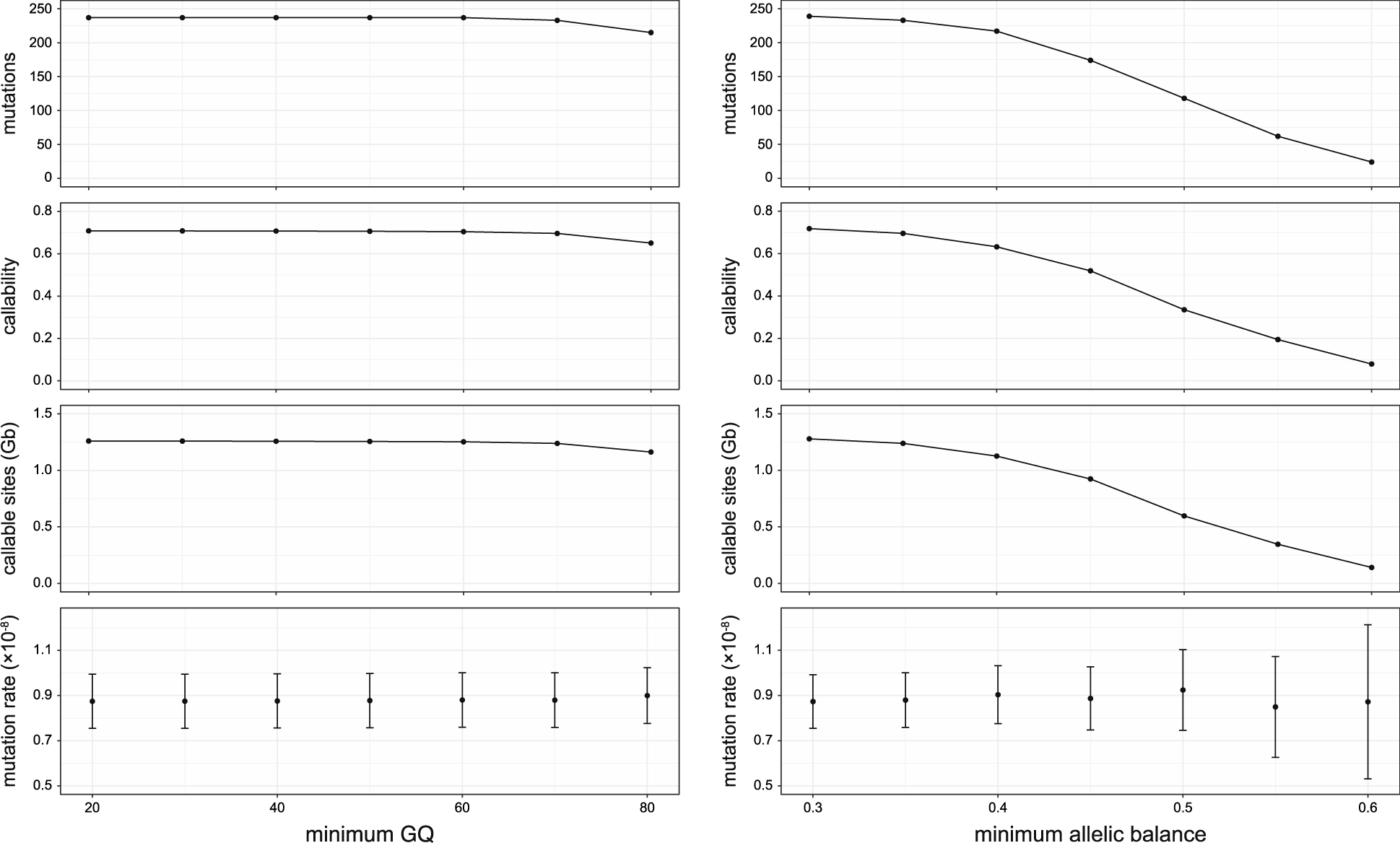
Mutation rates estimated with increasing filter stringency The number of mutations and the callable size of the genome decline with increasing filter stringency for (left) minimum genotype quality (GQ), while maintaining allelic balance > 0.35, and (right) minimum allelic balance, while GQ > 70. Error bars show 95% CI on estimates of the per generation, per bp rates under a Poisson model.

**Figure S3.**
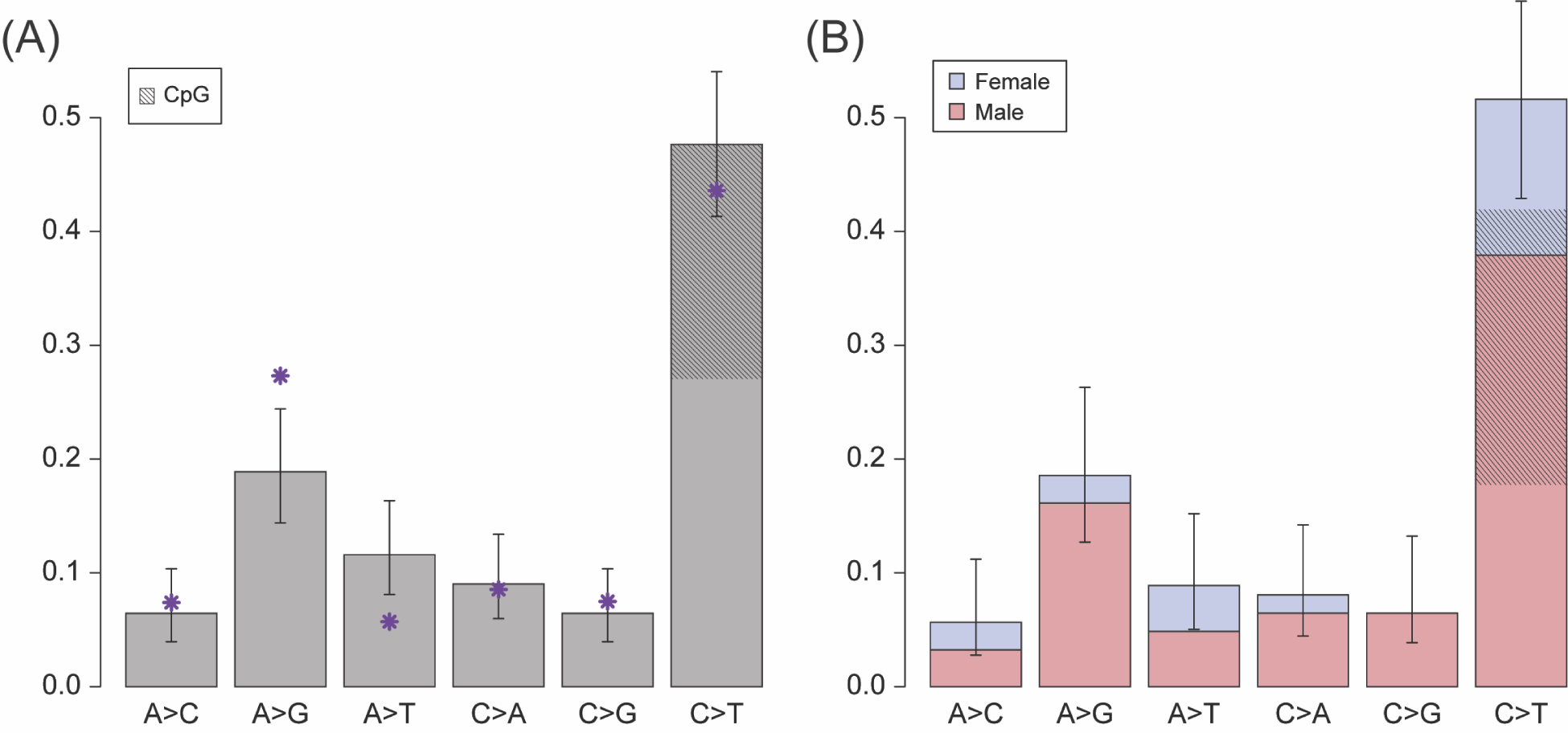
Cat mutation spectrum Proportion of each mutation class from mutations detected in all 11 trios. Asterisks (purple) show a comparison to the polymorphism spectrum for each mutation class (see Supplemental Results). (B) Proportion of each mutation class from only phased mutations. Shaded regions show the proportion of mutations occurring at CpG sites. 55% of phased C>T mutations occurred at CpG sites: 47% of these were of paternal origin and 8% were of maternal origin. Error bars show binomial 95% CI (Wilson score interval) for respective classes.

**Figure S4.**
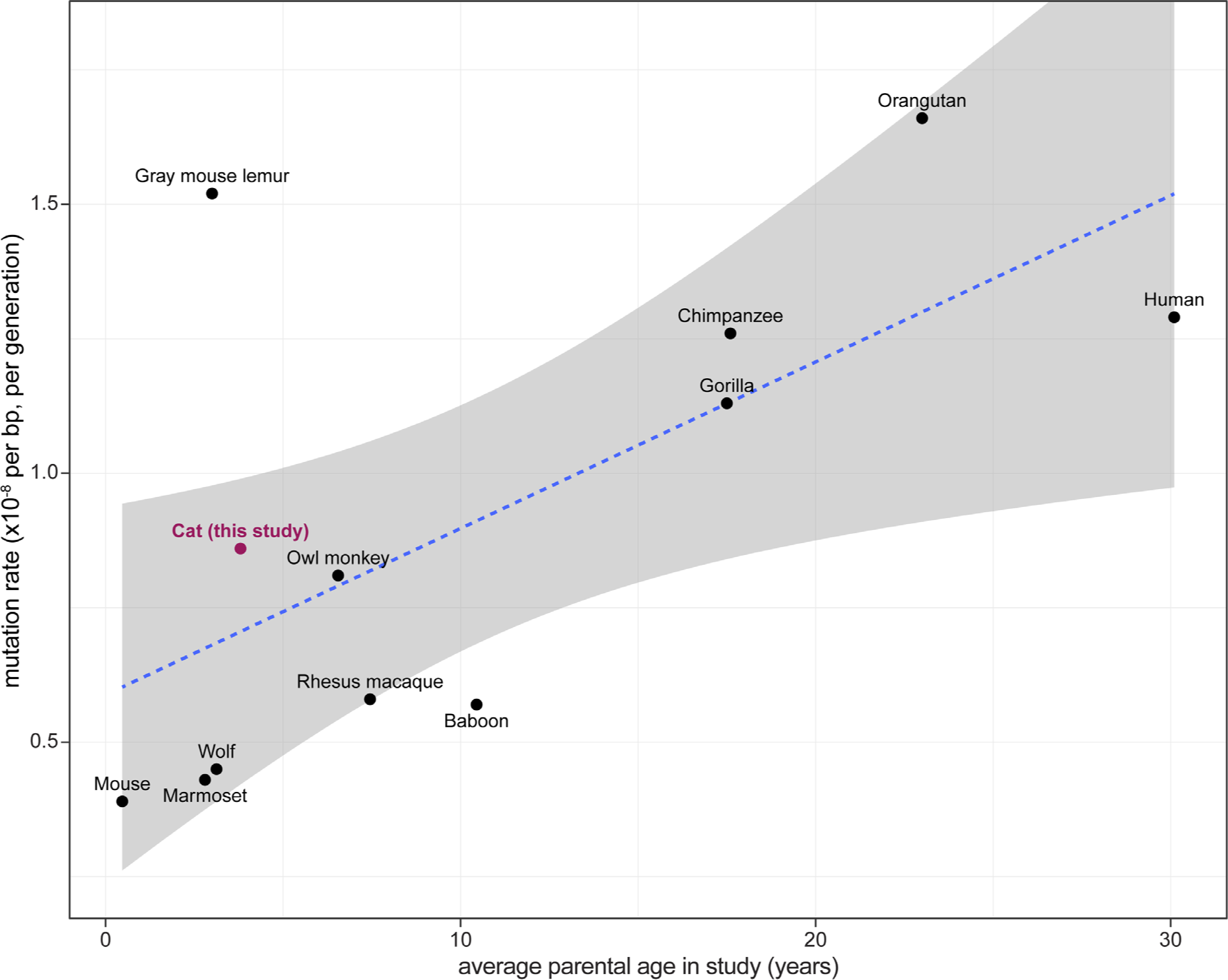
Mutation rate per generation and parental age in mammals Comparison of the per-generation mutation rate from the domestic cat (red) to rates from other pedigree studies in mammals. There is a significant relationship between the mutation rate and average parental age of animals used in each study (*p* = 0.02). Because these studies use different sampling approaches, we show a point estimate at the average age of animals used in each study to make the comparison. The blue dotted line shows a linear regression of the per-generation mutation rate with parental age from all presented estimates (95% CI in gray). References for each of these estimates: *Mouse*, Lindsay et al. (2019); *Wolf*, Koch et al. (2019); *Marmoset*, Yang et al. (2021); *Gray mouse lemur*, Campbell et al. (2021); *Owl monkey*, Thomas et al. (2018); *Rhesus macaque*, Wang et al. (2020); *Baboon*, Wu et al. (2020); *Chimpanzee*, Besenbacher et al. (2019); *Gorilla*, Besenbacher et al. (2019); *Orangutan*, Besenbacher et al. (2019); *Human*, Jónsson et al. (2017).

